# PRMT1 Modulates Alternative Splicing to Enhance HPV18 mRNA Stability and Promote the Establishment of Infection

**DOI:** 10.1101/2024.09.26.614592

**Authors:** David E.J. Williams, Kelly King, Robert Jackson, Franziska Kuehner, Christina Arnoldy, Jaclyn N. Marroquin, Isabelle Tobey, Amy Banka, Sofia Ragonese, Koenraad Van Doorslaer

## Abstract

Only persistent HPV infections lead to the development of cancer. Thus, understanding the virus-host interplay that influences the establishment of viral infection has important implications for HPV biology and human cancers. The ability of papillomaviruses to establish in cells requires the strict temporal regulation of viral gene expression in sync with cellular differentiation. This control primarily happens at the level of RNA splicing and polyadenylation. However, the details of how this spatio-temporal regulation is achieved still need to be fully understood.

Until recently, it has been challenging to study the early events of the HPV lifecycle following infection. We used a single-cell genomics approach to identify cellular factors involved in viral infection and establishment. We identify protein arginine N-methyltransferase 1 (PRMT1) as an important factor in viral infection of primary human cervical cells. PRMT1 is the main cellular enzyme responsible for asymmetric dimethylation of cellular proteins. PRMT1 is an enzyme responsible for catalyzing the methylation of arginine residues on various proteins, which influences processes such as RNA processing, transcriptional regulation, and signal transduction. In this study, we show that HPV18 infection leads to increased PRMT1 levels across the viral lifecycle. PRMT1 is critical for the establishment of a persistent infection in primary cells. Mechanistically, PRMT1 inhibition leads to a highly dysregulated viral splicing pattern. Specifically, reduced PRMT1 activity leads to intron retention and a change in the E6 and E7 expression ratio. In the absence of PRMT1, viral transcripts are destabilized and subject to degradation via the nonsense-mediated decay (NMD) pathway. These findings highlight PRMT1 as a critical regulator of the HPV18 lifecycle, particularly in RNA processing, and position it as a potential therapeutic target for persistent HPV18 infections.

## Introduction

Human papillomaviruses (HPVs) are small DNA viruses that infect cutaneous and mucosal epithelium and are among the most common sexually transmitted infections^1^. Long-term persistent HPV infections lead to 5% of all cancers worldwide, including nearly 250,000 cervical cancer cases each year, and are also associated with a variety of other anogenital and nasopharyngeal cancers^2,3^. These viruses are dependent on host cell machinery for replicating their DNA genomes. These infections induce cellular proliferation while delaying differentiation of keratinocytes within the infected epithelium; this is the basis for their oncogenicity^4^.

The viral lifecycle can be divided into three main phases. Following the initial infection of basal keratinocytes, the expression of viral genes (E1, E2, E5, E6, and E7) enables the virus to establish a productive infection^5^. Following *establishment*, the virus needs to be maintained in these actively dividing cells (i.e., *persist*). E1 and E2 proteins are necessary for the persistence and replication of the viral DNA^6,7^. The oncogenes E6 and E7 are highly associated with cellular transformation and carcinogenesis: e.g., E6 antagonizes p53 and promotes telomerase activity, while E7 antagonizes the tumor suppressor pRb^8,9^. Expression of these viral proteins plays critical roles in both establishment of the infection and the ability of the viral DNA to persist in dividing cells. These persisting infections are the leading risk factor for the progression of cancer^10,11^. Finally, to complete the viral lifecycle, cellular differentiation promotes a switch towards productive viral replication, packaging, and eventual virion release. In line with these lifecycle stages, viral gene expression of high-risk *Alphapapillomavirus* genomes progresses along three distinct expression programs: early (E6, E7, E1, E2, and E5), a combination of early and late (E1, E2, and E5), and exclusively late genes (L1 and L2). Initially, the early promoter, p105 for HPV18, drives the transcription of mRNAs encoding all early genes, which are polyadenylated at the early polyadenylation signal (pAE). These polycistronic mRNAs, which are composed of exons and introns, are subjected to alternative splicing involving specific splice sites. Therefore, mRNAs expressed from p105 have the potential to express E6, E7, E1, E2, E1^E4, and E5. However, the exact coding potential is decided through alternative splicing.

In sync with cellular differentiation, a shift occurs with the activation of the late promoter. This promotes high expression of mRNAs containing E1, E2, and E5 ORFs, omitting E6 and E7. Following this, E1 and E2 proteins facilitate viral DNA replication by binding to the origin of replication (ori) in the genome. As host cell differentiation advances, decreased activity of the early polyadenylation signal leads to read-through into the genuine late genomic region encoding L1 and L2. This results in polyadenylation at the late polyadenylation signal, yielding L2 mRNAs. Activation of specific late splice sites additionally generates L1 mRNAs alongside L2 mRNA. While promoter and poly-A activity regulate which viral mRNAs are expressed, extensive use of alternative splicing fine-tune viral gene expression^12^. It has been shown that alternative splicing and polyadenylation of HPV mRNAs is tightly regulated by positive and negative cis-acting RNA elements that interact with cellular serine–arginine-rich (SR) proteins and/or heterogeneous nuclear ribonucleoproteins (hnRNPs)^13–15^. Regulation of the viral transcriptome and by extension coding potential, could have important implications for the viral lifecycle. To understand how HPV18 gene expression is regulated during the viral life cycle we need to understand how HPV alternative splicing is controlled.

We performed a single-cell genomics approach to identify cellular factors correlated with viral gene expression shortly after infection (Williams et al., 2024). We further explored these expression data looking for host proteins that may regulate (viral) mRNA splicing. We identified an important role for protein arginine methyl transferase 1, PRMT1.

Protein arginine methylation is a post-translational modification specific to arginine residues. In mammals, nine methyltransferases are divided into type I and type II enzymes, according to their end products^16–20^. Type II PRMTs catalyze symmetric dimethylated arginines, while type I PRMTs lead to asymmetric dimethylarginine, the more common modification. Type I PRMTs (PRMT1, −3, −4, −6, and −8) seem to be more or less ubiquitously expressed^16,19^. These modifications play critical roles in diverse cellular processes like DNA repair^21–27^, RNA transcription^28–31^ and splicing^32–38^, and cellular differentiation^39–41^. PRMT1 is the predominant protein arginine methyltransferase in cells^42^.

In this manuscript we demonstrate that HPV18 upregulates PRMT1 expression throughout the viral lifecycle. Inhibition of PRMT1 impacts the ability of HPV18 genomes to establish a long-term infection in primary cells. Mechanistically, we demonstrate that inhibition of PRMT1 activity results in alternative splicing of HPV18 mRNA leading to a reduction in the steady-state levels of specific viral mRNAs. Finally, we demonstrate a role for nonsense mediated decay in regulating the viral transcript levels.

## Results

### HPV18 increases PRMT1 levels across the viral lifecycle

We previously used scRNA-Seq to identify differentially expressed genes between HPV18(+) and HPV-negative control cell lines. Briefly, cells were infected with quasiviruses and 10 days post-infection, cells were processed for 10X genomics single cells sequencing. Using this approach, we identified PRMT1 as a differentially expressed gene between HPV18(+) and parental control cells (Williams et al., 2025). Based on the scRNA-Seq data, HPV18(+) cells express higher levels of PRMT1 mRNA compared to mock infected cells at 10 days post infection (**Figure 1A**).

**Figure 1.**
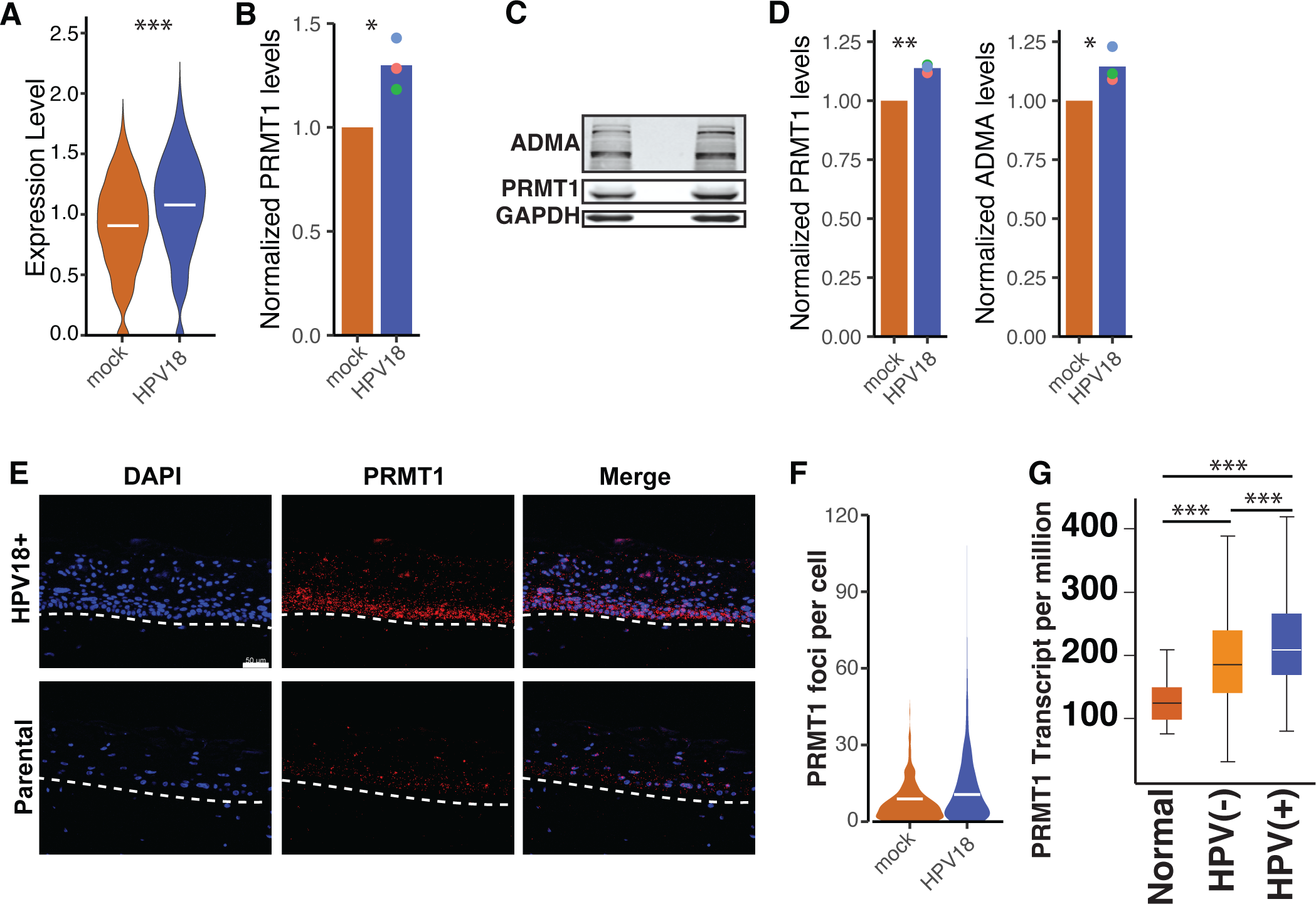
HPV18 upregulates PRMT1 levels throughout the viral lifecycle. (A) Violin plots showing the expression of PRMT1 based on scRNA-seq. Expression level is shown as normalized expression levels. (B) mRNA was isolated from matching parental and HPV18(+) primary cervical cells was amplified using qPCR. Colored dots indicate individual replicates, bars indicate the mean levels. Expression was normalized to parental cells. (C) Total protein was isolated from matching parental and HPV18(+) primary cervical cells and analyzed using western blot. Blots were probed for PRMT1 and Asymmetric dimethylated arginine (ADMA) levels. Protein levels were quantified in (D) and (E) respectively. Colored dots indicate individual replicates, bars indicate the mean levels. (F) Matching parental and HPV18(+) primary cervical cells were cultured as 3D organotypic rafts. PRMT1 mRNA was detected using RNAScope and (G) quantified according to the manufacturer’s guidelines. (F) PRMT1 expression levels were extracted from the TCGA using the UALCAN portal. Head and neck tumors were stratified by HPV. *p<0.05, **p<0.01, ***p<0.001.

Following infection, HPV18 establishes a long-term persistent infection. We mimic this part of the viral lifecycle by culturing HPV18(+) cells in the presence of terminally irradiated J2 fibroblasts^43^. As was seen 10 days post-infection (**Figure 1A**), HPV18 increases the steady state levels of PRMT1 in these stable cells at the mRNA (**Figure 1B)** and protein (**Figure 1C**) levels. Primary human cervical keratinocytes (HCKs) stably maintaining the viral genome had increased levels of PRMT1 protein and a resulting increase in proteome wide Asymmetric DiMethylated Arginine levels (ADMA). (**Figure 1C and quantified in 1D**). Taken together these data indicate that persistently infected keratinocytes upregulate PRMT1 levels (mRNA and protein) leading to an increase in cellular ADMA levels.

Papillomaviruses complete their lifecycle in stratified epithelia. To study this part of the viral lifecycle, we differentiated primary HPV18(+) and matched control cells using the described 3D organotypic raft culture. We used RNAscope to detect the expression of PRMT1 in these cells. In the control tissue, PRMT1 protein is most pronounced in the basal and suprabasal layers (**Figure 1E**). Similar to the 2D cell culture experiments, HPV18 (+) cells increase the expression of PRMT1 (**Figure 1E and quantified in 1F**). This indicates that HPV18 upregulates PRMT1 expression as the virus completes its lifecycle in 3D epithelia.

Finally, long term-persistent infections can drive transformation of the host cell. To investigate whether HPV upregulates PRMT1 during cancer progression, we analyzed available head and neck cancer data from the TCGA^44^. Unlike cervical cancers, head and neck cancers can be both HPV(+) and HPV(-) allowing us to identify a potential role for HPV infection. Based on the TCGA data, **Figure 1G** shows that while PRMT1 is upregulated in HPV(-) cancers, PRMT1 levels are further upregulated in HPV(+) cancers (blue bar).

In conclusion, HPV upregulates PRMT1 shortly after infection and during persistent infection. Furthermore, HPV18 changes the spatio-temporal expression of PRMT1 during tissue differentiation, suggesting that PRMT1 may be an important host factor for an optimal viral lifecycle. Finally, PRMT1 levels are further increased in HPV(+) head and neck cancers.

### Loss of PRMT1 activity reduces steady state levels of canonical viral transcripts in the cell

The above data demonstrate that HPV18 upregulates PRMT1 levels and associated ADMA levels in the infected cell, suggesting that PRMT1 may have a pro-viral function.

To test this experimentally, we treated persistently infected HPV18(+) HCKs with 2 μM GSK-3368715 (a selective PRMT1 inhibitor; PRMT-i) for 3 days. Treatment with PRMT-i reduced proteome wide ADMA levels (**Figure 2A** and **2B**). At the mRNA level, inhibition of PRMT1 led to a previously described reduction in FOXM1 (**Figure 2C**; light grey) transcripts. Importantly, while expression of other host genes (e.g., PTPB2. **Figure 2C**; dark grey) was not reduced. inhibition of PRMT1 significantly reduced the steady state levels of the E6*I (yellow), E1^E4 (green), and E5 open reading frames (**Figure 2C**). This reduction in HPV18 mRNA was matched by a concordant decrease in E7 protein levels (**Figure 2D** and **2E**).

**Figure 2.**
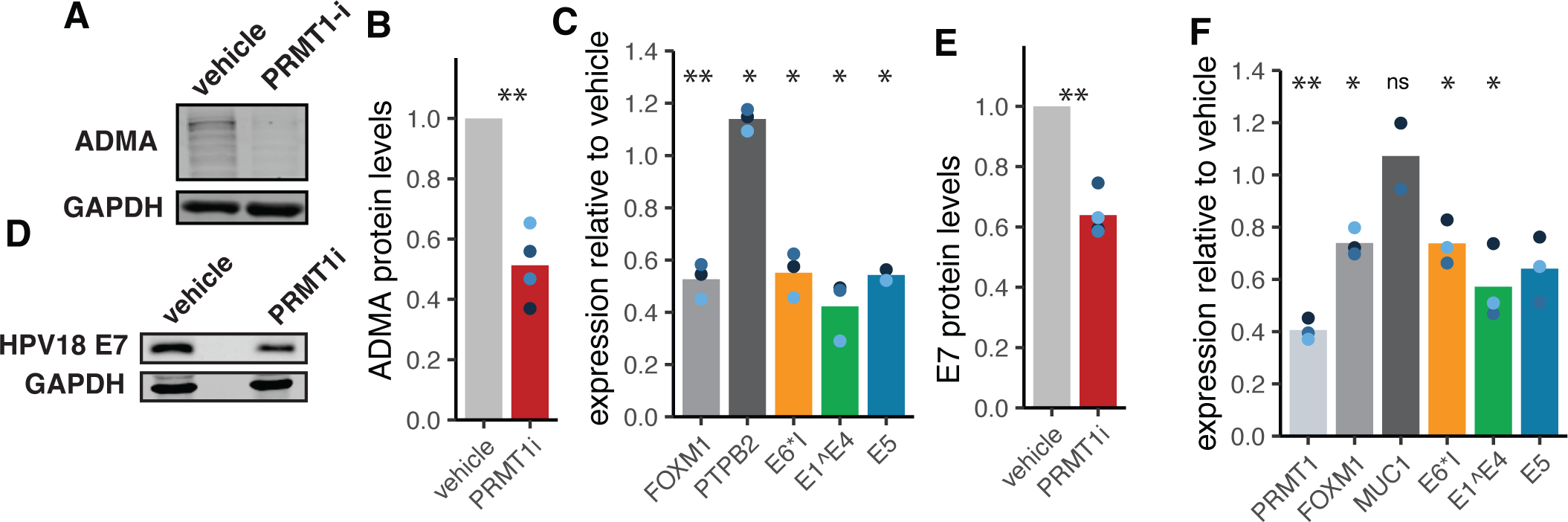
PRMT1 is essential for optimal processing of viral mRNA. (A) Primary cervical cells were treated with 2uM PRMT1i for 3 days. Total protein was isolated from matching parental and HPV18(+) primary cervical cells and analyzed using western blot. Blots were probed for Asymmetric dimethylated arginine (ADMA) levels and quantified in (B). (C) Lab derived HPV18(+) primary cervical cells were treated with treated with 2uM PRMT1i for 3 days mRNA was isolated from vehicle and inhibitor treated cells and amplified using qPCR. (D) Protein was isolated from cells treated as in (C) and probed for HPV18 E7 levels and quantified in (E). (F) Primary cervical cells were treated with doxycycline for 3 days and infected using HPV18 quasivirus. Four days post infection, mRNA was isolated from vehicle and doxycycline treated cells and amplified using qPCR. Throughout, colored dots indicate individual replicates, bars indicate the mean levels. *p<0.05, **p<0.01, ***p<0.001.

To further establish a role of PRMT1 on viral transcriptional regulation following infection, we generated primary human cervical keratinocytes expressing a doxycycline inducible shRNA targeting PRMT1. Cells were treated with doxycycline to induce the expression of shRNA and the knockdown of PRMT1. Three days post-induction, cells were infected with HPV18 quasiviruses and total mRNA was isolated 4 days post infection. We titrated doxycycline to achieve a roughly 60% knockdown of PRMT1 mRNA (**Figure 2F; light grey bar**). Furthermore, this knockdown reduces the expression of FOXM1, a published PRMT1 target gene^41^, without altering the expression of MUC 1, a cellular control gene (**Figure 2F; grey and dark grey bar, respectively)**. We quantified the viral mRNA 4 days post infection and observed a significant reduction in steady state HPV18 mRNA. Importantly, E6*I (yellow), E1^E4 (green), and E5 (blue) levels are reduced to equivalent levels as FOXM1 levels (**Figure 2F**), further supporting the idea that PRMT1 regulates the cellular levels of these viral transcripts. Overall, these data indicate that PRMT1 is essential for optimal regulation of viral transcript levels.

### PRMT1 is essential for the efficient establishment of a persistent HPV infection

Given the observation that PRMT1 is upregulated at each stage of the viral lifecycle (**Figure 1**), and PRMT1 function is essential for viral mRNA levels both shortly after infection and during persistent infection (**Figure 2**), we hypothesize that PRMT1 may be needed for optimal establishment and persistence of the viral genome. We used a previously described establishment assay in which the late region of HPV18 is replaced with a G418 resistance expression cassette. Following infection, G418 selects cells that maintain the recombinant viral genome^43^. Importantly, this approach does not select for integration of the viral genome into the host cell but allows for extra-chromosomal maintenance of the HPV18 genome^43^.

As described in **Figure 3A**, cells were treated with doxycycline to induce the expression of PRMT1 shRNA 3 days before infection and maintained shRNA mediated knockdown for a total of 10 days (i.e., seven days post infection). G418 selection was started 2 days after infection and was continued until macroscopically visible colonies could be observed on the vehicle treated plates. In parallel experiments, uninfected cells were grown in the presence of doxycycline (or vehicle) for 10 days but without G418 selection. The population doublings of these cells were tracked (**Figure 3B**) and no differences in growth was observed, suggesting that a 10-day knockdown of PRMT1 does not alter cellular replication.

**Figure 3.**
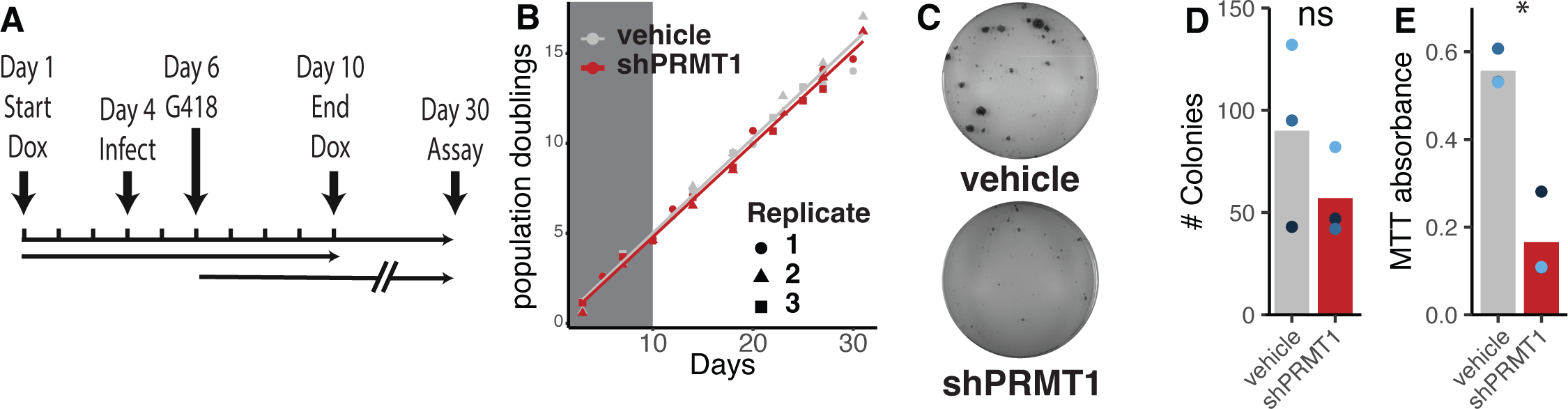
PRMT1 is essential for optimal establishment and maintenance of HPV18 infection. (A) Timeline of the experiment. (B) Parental, HPV(-) cervical cells were treated with doxycline (or vehicle control) to induce the expression of PRMT1 specific shRNA for 10 days (grey box). Cells were cultured for an additional 20 days and population doublings were tracked. (C) Primary cervical cells were treated with doxycycline for 3 days and infected using HPV18 quasivirus as in Figure 4. ∼30 days post infection, J2 fibroblast feeders were removed and cells were stained with MTT reagent. Gross images were captured and counted (D). (E) MTT was quantified and plotted. Throughout, colored dots indicate individual replicates, bars indicate the mean levels. *p<0.05, **p<0.01, ***p<0.001.

When macroscopic keratinocyte colonies were visible on the control plates, (roughly 30 days post infection) the J2-3T3 fibroblasts were selectively removed, and keratinocytes were stained with MTT and imaged for gross colony count (**Figure 3C** and **D**). After imaging, the level of MTT metabolized to MTT-formazan was quantified as a proxy for total cell number on each plate (**Figure 3D**). Infection of vehicle treated cells, expressing wildtype levels of PRMT1, supported robust establishment of infection, with numerous large colonies (**Figure 3C**). shRNA mediated knockdown of PRMT1 resulted in a 70% reduction in viable cells as quantified with the MTT assay (**Figure 3E**). However, there is no statistically significant difference in the ‘absolute’ number of colonies (**Figure 3D**). Taken together, this suggests that while cells with reduced PRMT1 are still susceptible to infection (and thus are initially resistant to G418) the viral genomes were lost as the cells divided resulting in non-amplifying colonies.

### PRMT1 does not regulate HPV18 p105 promoter activity

We demonstrate that reduction of PRMT1 (activity) leads to a decrease in steady-state levels of viral mRNA. We first asked whether PRMT1 regulates HPV18 promoter activity. The viral P105 promoter is located in the viral upstream regulatory region (URR). To evaluate viral promoter activity, we used a construct expressing firefly luciferase under the control of the entire HPV18 URR. We treated primary cells with 2 μM PRMT-i for 2 days and transfected cells with the HPV18_URR-firefly plasmid along with a Renilla expressing control plasmid^45^ (CMV promoter). Inhibition of PRMT1 leads to an increase in luciferase signal, indicating increased P105 activity (**Figure 4A**). We also used shRNA to knock-down PRMT1 levels as described above (**Figure 4B**) and 6 days post the start of knock-down, cells were transfected with both plasmids. One day after transfection, we quantified the luciferase signal as a proxy for viral promoter activity (**Figure 4C**). Therefore, while loss of PRMT1 (activity) reduces viral mRNA levels, it does not decrease (and may increase) viral promoter activity, suggesting that the observed differences in steady-state viral mRNA are likely regulated post-transcriptionally.

**Figure 4.**
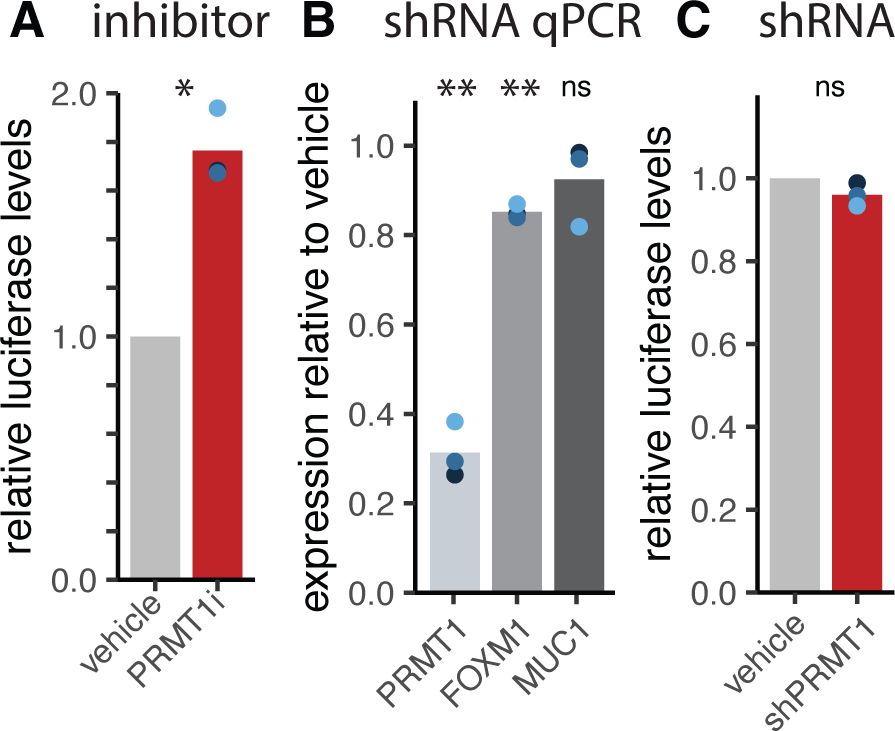
PRMT1 does not reduce HPV18 early promoter activity. (A) Primary cervical cells were treated with or 2uM PRMT1 inhibitor for 2 days two days and transfected using a firefly-luciferase reporter controlled by the HPV18 URR and a renilla-luciferase (CMV promoter) control plasmid. 24 hours post transfection, the relative luciferase levels were quantified and plotted. (B) Primary cervical cells were treated with 640 ng/ml doxycycline for six days days and transfected using the same plasmids as (A). mRNA was isolated 3 days after induction (i.e., at time of transfection) and quantified using qPCR. Luciferase activity was quantified as in (A). Throughout, colored dots indicate individual replicates, bars indicate the mean levels.

### PRMT1 promotes alternative splicing of viral mRNA

Given its known role in regulating cellular splicing, we hypothesized that PRMT1 may be regulating viral mRNA at the level of mRNA splicing. We treated primary foreskin derived cells stably containing HPV18 with 2 μM PRMT1i and extracted total RNA after 3 days of treatment. Viral mRNA was enriched using a custom HPV18 specific set of probes (see materials and methods). Enriched mRNA was sequenced at the University of Arizona Genomics Core. This experiment was performed four independent times (indicated by different colors). The sequencing reads were aligned to the HPV18 reference genome using STAR, a splice-aware aligner. Changes in viral splice junction usage were visualized using Sashimi plots (**Figure 5A**). The top (vehicle) and bottom (PRMT1i) plots show RNA-Seq read densities along the HPV18 early region. The arching lines show the top 10 observed exon-exon junctions (colored lines refer to qPCR amplicons used throughout this manuscript. These 10 junctions are quantified for each individual replicate (**Figure 5B)**. Focusing on the vehicle treated (top) coverage map, we detect the previously reported HPV18 transcript map. For example, there is a drop in coverage between the SD233 and SA416 corresponding to the splicing event giving rise to E6* (orange arch). Similarly, the splice junctions giving rise to E1^E4 (SD929 to SA3434; green arch) are readily visible. These cells primarily express mRNAs processed to express E6*, E7, E1^E4, and a relatively small amount of full length E6 and E2. Of note, we identify a prominent splice event between SD3706 and SA4159 (red arch) which would remove the C-terminal halve of E2 and the E5 ORF. To our knowledge this putative splice junction has not been previously described.

**Figure 5.**
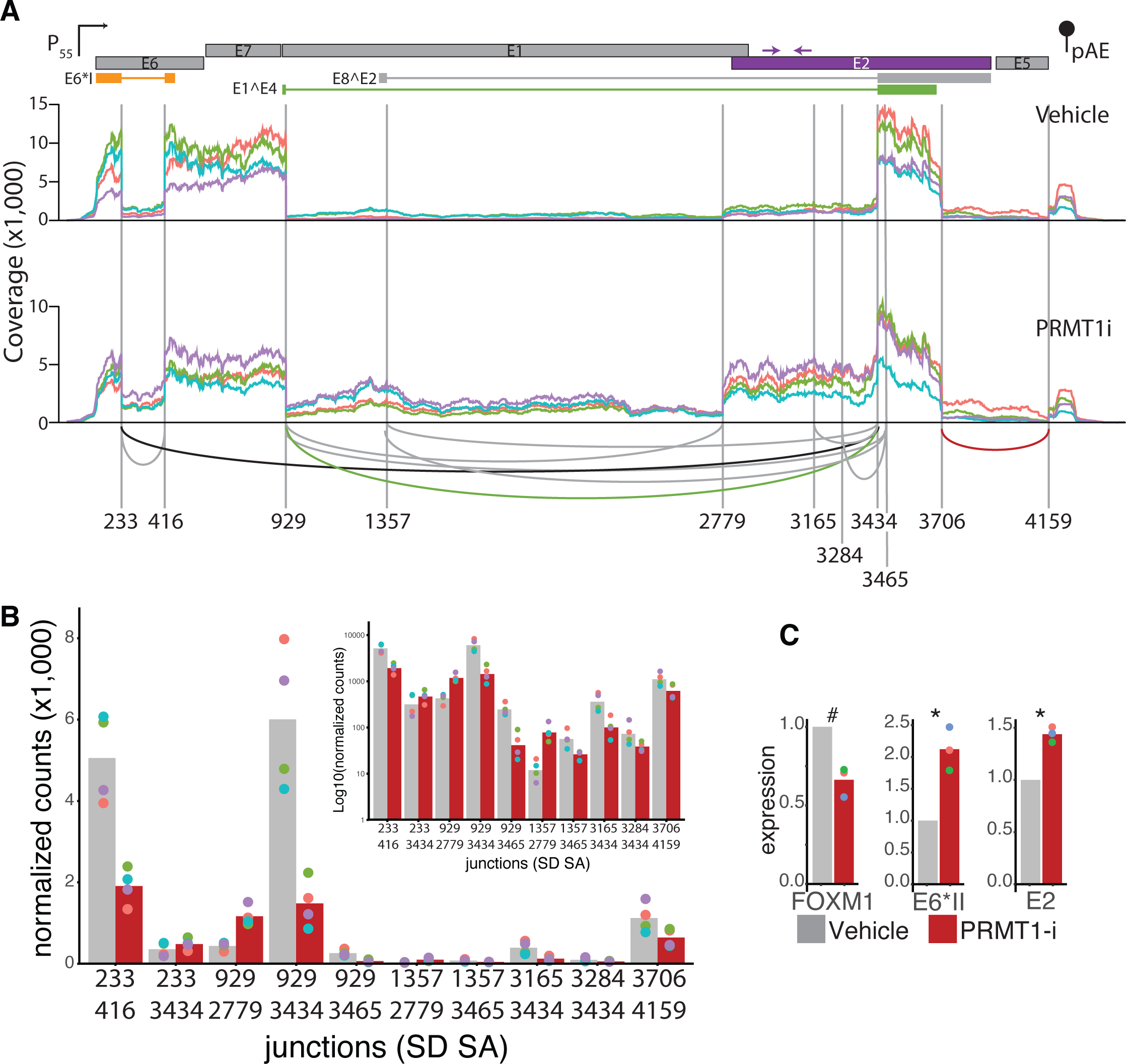
PRMT1 promotes alternative splicing of viral mRNA. (A) Lab derived HPV18(+) foreskin derived cells were treated with 2uM PRMT1i (or vehicle control) for three days. Viral mRNA was captured and sequenced. HPV18 coverage plots relative to a cartoon of the early region of the HPV18 genome. Colored lines indicate different biological replicates. The top graph is vehicle treated, while the bottom is representative of the viral transcriptome in PRMT1i treated cells. Arches connect the top 10 observed splice donor and acceptor sites. The colors of these arches are relative to the viral genes as used in other figures. Numbers indicate splice donors or acceptors. (B) Splice junctions are quantified as normalized counts (inset shows log10 to highlight lower expressed sites). (C) Lab derived HPV18(+) cervical cells were treated with 2uM PRMT1i (or vehicle control) for two days. mRNA was isolated and quantified using qPCR with the indicated primers.

The PRMT1i (bottom) coverage map shows that inhibition of PRMT1 reduces the overall number of reads mapping to HPV18 without altering which splice sites are combined (**Figure 5A**). However, the RNA-Seq read densities suggest that PRMT1 inhibition alters the ratios of which mRNA are produced. For example, inhibition of PRMT1, reduces the SD233 - SA416 junction by roughly 50% (compare grey to red bar in **Figure 5B**). Together this indicates that PRMT1 inhibition leads to a reduction in E6* splicing and a relative increase in the expression of mRNAs containing full length E6. Even more striking is the region between SD929 and SA3434. Inhibition of PRMT1 leads to an increase in sequence coverage in this region. Specifically, there is an increase in coverage from SD929 to SA2779 and a further increase from SD2779 to SA3434. Functionally, this is expected to result in a relative increase in the expression of the replication proteins E1 (nt. 914 to 2887) and E2 (nt. 2817 to 3914). No good E1 or E2 antibodies are available, so this was not further tested. This change in sequencing coverage can be explained through an increase in splicing between SD929 and SA2779 (**Figure 5B**). In parallel, there is a roughly 2-fold reduction in usage of the E1^E4 specific splice site (SD929 to SA3434 junction). Taken together, these analyses suggest that inhibition of PRMT1 leads to intron retention and an increase in viral transcripts that could express full length E6, E1, and E2. As the E7 protein is believed to be translated from E6* containing transcripts^46^, these data would help explain the observed reduction of the E7 protein (**Figure 2G).**

To confirm these RNASeq based results, we repeated the inhibitor treatment in a separately generated set of HPV18(+) cervical cells and analyzed mRNA expression using qPCR with different primers. The first set detects the E6*II transcript (SD 233 – SA 3444; black arch in **Figure 5A**), while the second set is specific to full length E2 and is located upstream of the overprinted E4 open reading frame (purple arrows in **Figure 5A**). Using these primers, we confirm an increase in both E6*II and full length E2 containing mRNA species following inhibition of PRMT1 **(Figure 5C)**.

These data demonstrates that PRMT inhibition leads to alternative splicing events in the viral mRNA.

### PRMT1 promotes intron exclusion on viral mRNA

While STAR quantifies splice junctions, we also wanted to perform a more thorough statistical comparison on predicted full length, spliced, mRNA species. We aligned the HPV18 reads to the reference transcripts^12^ using SALMON and further analyzed the expression of individual transcripts using DESeq2. **Figure 6A** shows a cartoon representation of the different mRNA species with differential expression shown on a log2 fold scale. Transcripts indicated by ** are statistically differentially expressed between PRMT1i and vehicle. **Figure 6B** shows the same data, but instead of showing fold changes, it shows the relative expression of each transcript. In Figure 6B, transcript counts are normalized by total reads mapped to HPV18, to allow for relative comparison between both conditions. For example, species K has the biggest log2 fold change, but it is relatively lowly expressed.

**Figure 6.**
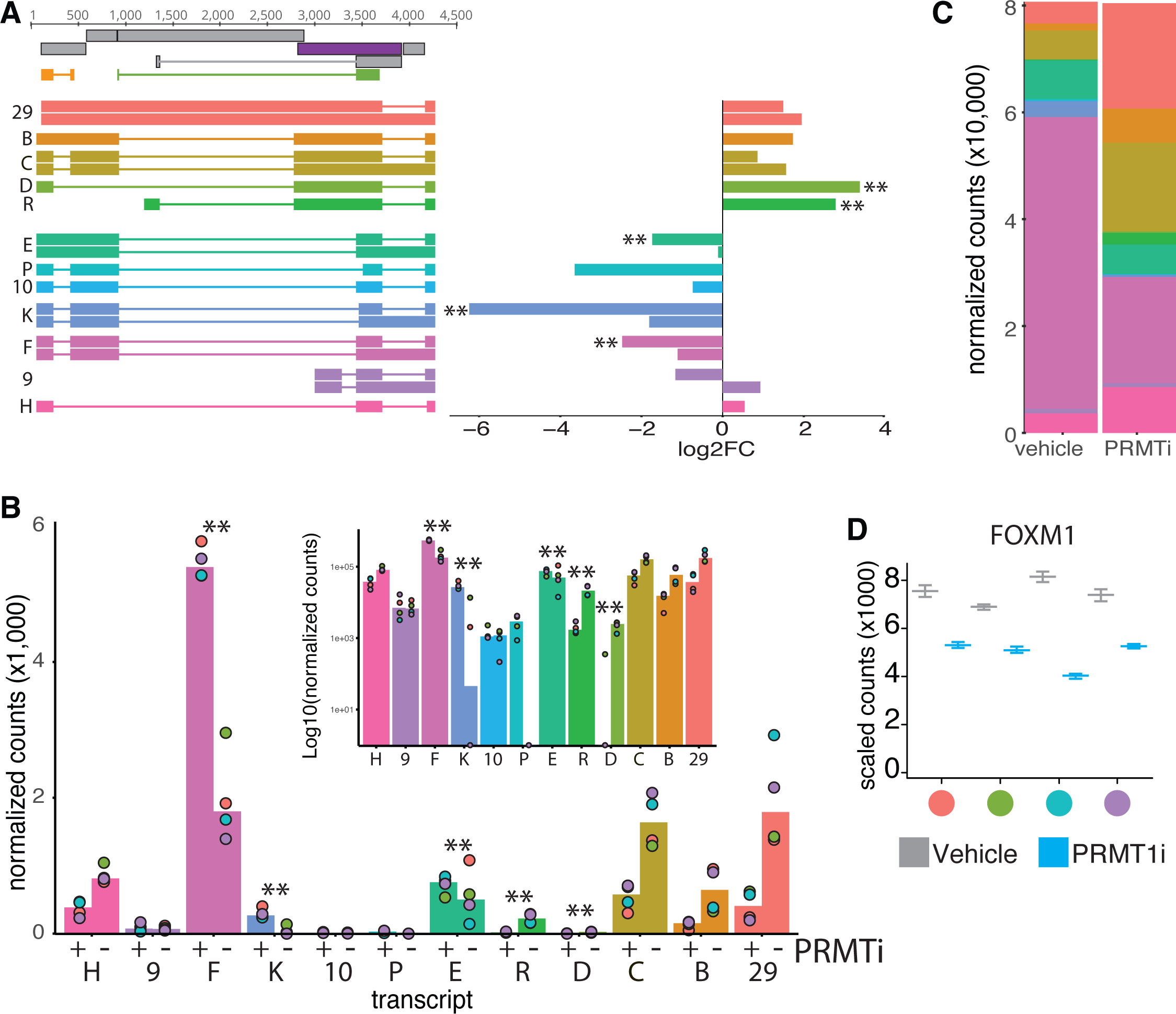
Loss of PRMT1 activity promotes intron retention. This figure is based on the same RNASeq data presented in Figure 5. (A) Cartoon representation of analyzed viral transcripts with log2 fold expression as quantified using SALMON. Transcripts with and without the SD3706-SA4159 (in E2-E5) splice are shown separately. Colors of the expression bars match the colors of the transcripts. (B) Transcripts are plotted as normalized counts (inset shows log10 to highlight lower expressed species). The x-axis matches the transcripts to the cartoon in (A). Transcripts with and without the SD3706-SA4159 (in E2-E5) splice are combined. (D) Quantification of FOXM1 levels in 4 biological replicates. Colors at the x-axis match the replicate colors in Figure 5 and 6.

This analysis identified 5 viral transcripts that are statistically (Log2 FC >2 and p<0.01) significantly differentially identified under both conditions (**Figure 6A**). Inhibition of PRMT1 statistically upregulates 2 viral transcripts (species D and R). These mRNA species utilize the splice acceptor at 2779 just upstream of the E2 start codon. Loss of PRMT1 activity leads to the (statistical) downregulation of three mRNA species (E, F, and K). Species F is the most abundant transcript in the vehicle treated cells and uses the canonical E6*I and E1^E4 associated splice sites (SD233-SA416 and SD929-SA3434, respectively). Species E and K also splice E6*I and utilize the canonical E1^E4 SD929 splice-donor, but splice into non-canonical splice acceptors within E2.

It is hard to define introns and exons for papillomavirus genes as almost every part of the genome encodes for protein. Therefore, to further investigate the idea of intron inclusion, we counted the number of potential splice donor-acceptor pairs in each transcript. In **Figure 6C** we sorted the transcripts by the number of potential splice donor-acceptor pairs, with the fewest remaining pairs on the bottom (species H) and the most pairs on the top (species 29). The size of the graph corresponds to the normalized counts for each transcript as in Figure 6B. Overall, this graph shows that PRMT1 inhibition causes a reduction in the most spliced (i.e., fewer potential SJs left) forms and an increase towards the larger species. This analysis further supports the idea that PRMT1 regulates intron removal from viral mRNA.

To confirm the activity of the PRMT1i, we quantified the FOXM1 expression levels from these RNASeq samples. PRMT1i treatment resulted in a roughly 40% decrease in FOXM1 levels, in line with other figures in this manuscript. The colored dots on the x-axis represent the biological replicates and the colors match the replicates throughout **Figures 7** and **8**.

**Figure 7.**
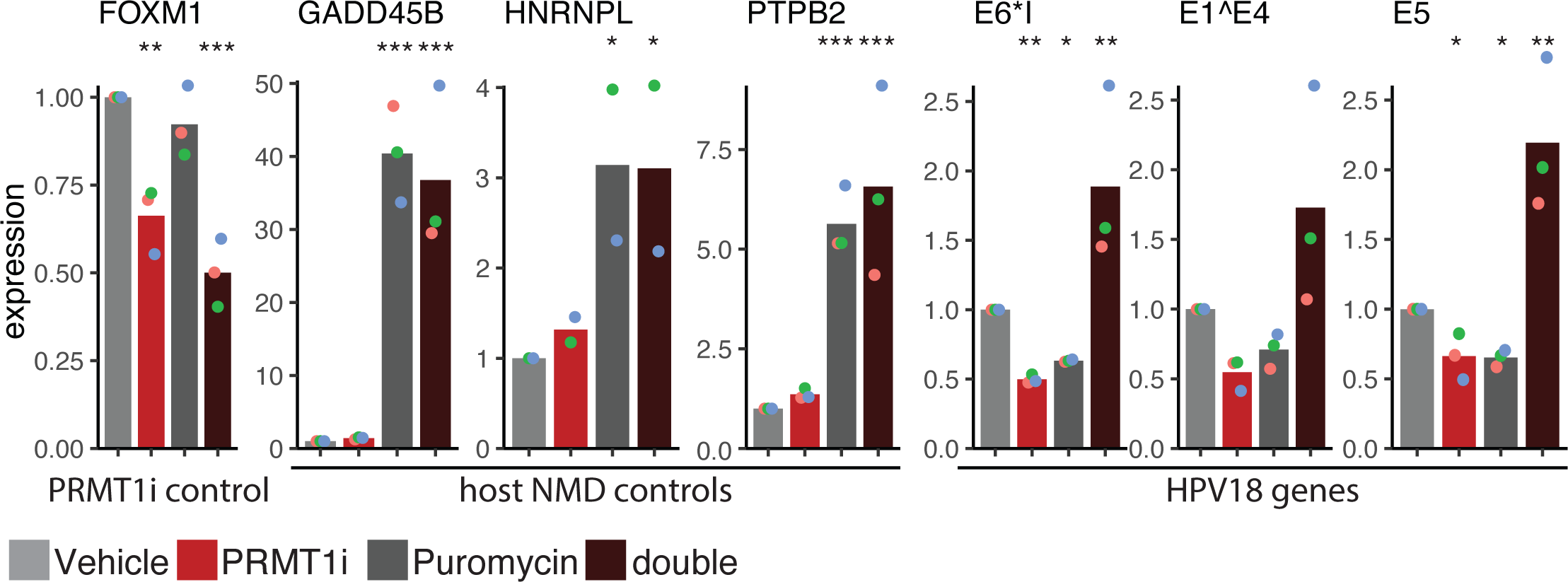
Inhibition of PRMT1 leads to alternative splicing coupled nonsense mediated decay. (C) Lab derived HPV18(+) foreskin derived cells were treated with 2uM PRMT1i (or vehicle control) for 3 days. Cells were treated with puromycin for an additional 8 hours. Transcript levels were quantified for FOXM1, GADD45B, HNRNPL, and PTPB2 (NMD controls), and three viral transcripts. Throughout, colored dots indicate individual replicates, bars indicate the mean levels.

### Loss of PRMT1 destabilizes viral transcripts through nonsense mediated decay

We demonstrate that PRMT1 is essential for regulated splicing (**Figures 7** and **8**) and maintenance of steady-state levels (**Figure 2**) of specific viral mRNA.

A potential mechanism for PRMT1 mediated stability of HPV mRNA is alternative splicing coupled Nonsense Mediated Decay^47^. To test a role of Nonsense Mediated Decay in regulating HPV18 mRNA levels we treated HPV18(+) HCKs with 2uM PRMT1 inhibitor or vehicle control for 40 hours. At this point, cells were treated with puromycin (or control) to inhibit nonsense mediated decay for another 8 hours (**Figure 8**). Total RNA was extracted and analyzed by qPCR for viral and host transcripts. As before, PRMT1 inhibition reduces the levels of FOXM1 mRNA in these cells. Inhibition of nonsense mediated decay did not impact FOXM1 levels, while the combination treatment reduced cellular FOXM1 mRNA levels, demonstrating that FOXM1 is regulated by PRMT1 in a nonsense mediated decay independent manner as reported. To ensure that puromycin blocked nonsense mediated decay, we included primers for GADD45B, HNRNPL, and exon 2 of PTB2. These mRNAs are known to be regulated by nonsense mediated decay^48,49^. As previously reported, treatment with puromycin resulted in a 30-fold increase in GADD45B mRNA, 3 fold increase in HNRNPL, and a 5-fold increase of the PTPB Exon 2 containing mRNA compared to the vehicle only sample (**Figure 8**). PRMT1 inhibition did not impact the levels of these host transcripts. These data indicate that puromycin treatment blocks nonsense mediated decay in our system and that PRMT1 inhibition does not typically result in changes to nonsense mediated decay.

As before, PRMT1i treatment alone resulted in a decrease in E1^E4, E6*I, and E5 mRNA levels. The additional inhibition of NMD leads to a ∼4-fold increase in viral mRNA (compare red to black). Taken together, these data suggest that PRMT1 controls viral mRNA steady state levels by coupling of alternative splicing and nonsense-mediated mRNA decay. Interestingly, Inhibition of nonsense mediated decay also reduced the levels of these viral transcripts. While we do not totally understand this phenotype of nonsense mediated decay inhibition leading to a reduction in target gene expression was previously seen for MMP9^50^.

## Discussion

In this study we show that HPV upregulates the expression of PRMT1 mRNA, protein, and enzymatic activity at every stage of the viral lifecycle: from infection, to productive replication, and during oncogenic progression. Furthermore, we demonstrate that inhibition of PRMT1, either by shRNA or small molecule inhibitors reduces the ability of HPV18 to establish a persistent infection in primary cervical keratinocytes.

PRMT1 has been demonstrated to be necessary for the replication and transcriptional activation of diverse DNA viruses. PRMT1 regulates Herpes Simplex replication through methylation of the HSV1 ICP27 protein^51^. Methylation of the KSHV LANA protein modulates a wide array of LANA functions during latency^52^. PRMT1 binding to the EBNA1 protein from EBV is vital for viral replication and segregation to daughter cells^53^, critical functions involved in the long-term persistence of these viruses. PRMT1 overexpression reduces the transcription of hepatitis B virus (HBV) in a methyltransferase dependent manner^54^. Efficient Adenovirus type 5 virion production depends on the methylation of one of the viral proteins by PRMT1^55^. Together, these data demonstrate that PRMT1 plays important roles throughout the lifecycle of diverse DNA viruses. Furthermore, viruses have been shown to alter PRMT1 activity in cells. The HBV X protein, HBx, inhibits PRMT1 activity^54^, while the EBV LMP1 protein induces PRMT1 expression^56^.

In our study, we provide compelling evidence that inhibition of PRMT1 causes intron retention leading to nonsense mediated decay of a subset of viral transcripts. Viral transcription is carefully regulated to allow for spatio-temporal expression of viral proteins. By inhibiting PRMT1, this careful regulation is disturbed by changing the ratio of viral transcripts. Regulation of the papillomavirus transcriptome is not fully understood. From our data, it is clear that PRMT1 regulates intron retention and exclusion and that this plays important roles in regulating the viral lifecycle. However, due to the small viral genome, the distinction between intron and exon is complicated and transcript dependent; an intron on one transcript is a protein coding exon on another transcript. The implications of this regulation remain to be determined. Specifically, it has been demonstrated that E7 is expressed from transcript in which the E6 intron is removed^46^. Therefore, regulation of the E6*I splice site regulates the amount of E6 and E7 oncogene expression. The inhibition of PRMT1 leads to intron retention and increases the expression of full length E6 while reducing the expression of E7. It is a believed that a careful balance of both oncoproteins is essential for an optimal viral lifecycle^8,9^. It is likely that similar regulation of viral translation exists for other proteins, but this remains to be determined.

It is unclear why papillomaviruses evolved this intricate regulation of viral gene expression. However, there is some evidence suggesting that control at the transcript/splicing level provides an evolutionary benefit under certain conditions. As mentioned above, E6 and E7 are produced from separate transcripts due to alternative splicing of the E6* intron/exon. While this is the case for the so-called high-risk papillomavirus genomes, the other members of the genus *Alphapapillomavirus,* use two distinct promoters to transcribe E6 and E7^57^. It is tempting to speculate that alternative splicing allowed for an additional level of post-transcriptional fine-tuning of the viral proteins. Specifically, when combined with nonsense mediate decay, alternative splicing would allow for fine tuning of the viral transcriptome that may not be achievable simply through promoter regulation.

Alternative splicing is regulated through the action of RNA binding proteins. HPV) require constitutive and alternative splicing to generate mRNAs encoding the many essential proteins that are required to initiate, maintain and complete their life cycles. Alternative splicing dramatically expands the coding potential of papillomaviruses^57–69^. As with host mRNA, viral RNA splicing is regulated through RNA binding protein. For example, SR and hnRNP proteins have been shown to control viral RNA processing during infection^70–72^. In turn, RNA binding proteins are targeted by protein arginine methyltransferases (PRMTs) to regulate RNA splicing^20,65,73–79^.

Our data propose a role for nonsense mediated decay in regulating the viral transcriptome. Nonsense mediated decay is a post-transcriptional RNA quality control mechanism. Traditionally, nonsense mediate decay is believed to detect transcripts with in-frame premature termination codons and target these faulty mRNAs for degradation. While mutations or alternative splicing events can cause these pre-mature stop codons, it is becoming evident that alternative splicing coupled nonsense mediated decay (AS-NMD) is an important mechanism to regulate gene expression. Specifically, by altering the splicing towards the expression of a non-functional NMD-targeted isoform of the active gene, the effective translation of the protein is reduced^47,80–83^. Furthermore, in addition to premature stop codons, the length of the 3’-UTR and/or the presence of upstream open reading frames, can impact AS-NMD^84^.

Nonsense mediated decay restricts virus replication in both plant and animal cells. It is therefore not surprising that viruses evolved ways to counteract NMD^85^. As reviewed^85^, viruses can both encode an anti-NMD element in their genome or express a protein that counteracts NMD. For example, the positive-sense RNA retrovirus Rous sarcoma virus (RSV), encodes an an RNA stability element (RSE) to avoid NMD^86^. Zika virus (ZIKV) infection attenuates NMD. Mechanistically, the ZIKV capsid protein physically associates with both UPF3X and UPF1, leading to proteasomal degradation of UPF1 and blunting NMD^87,88^. Similar to HPV, Kaposi’s sarcoma-associated herpesvirus (KSHV/HHV8) persists as a viral episome in infected cells. Alternative splicing of KSHV mRNA generates NMD targets. Indeed, inhibition of NMD triggers increased lytic reactivation of KSHV in cellular models, indicating that NMD restricts lytic reactivation. Interestingly, one of the mRNAs targeted by NMD encodes the replication and transcription activator (RTA) protein, which is essential for RTA is required for KSHV reactivation. These data suggest that KSHV may subvert NMD to regulate the different phases of the viral lifecycle. It is not clear if whether KSHV utilizes mechanisms to subvert NMD remains to be examined^89^. Based on these observations, we speculate that HPV18 may use PRMT1 to regulate viral gene expression and avoid expressing NMD targeted viral mRNAs.

In this paper, we focused on how PRMT1 impacts the infection, establishment and persistence phase of the viral lifecycle. However, throughout their lifecycle, HPVs manipulate and depend on cellular differentiation. We show that HPV18 upregulates the expression of PRMT1 protein in 3D tissues. In differentiated cells, a differentiation-inducible ‘late’ promoter is derepressed^90^, leading to the expression of the L1 and L2 genes. Given the role of PRMT1 in regulating the early viral transcriptome, it is possible that maintained expression of PRMT1 in the differentiated tissues also regulates the late transcriptome.

In conclusion, we demonstrate through genomics and cell biology that PRMT1 is essential for an optimal viral lifecycle, in part, due to a role of PRMT1 in regulating the viral transcriptome through alternative splicing and nonsense mediated decay.

## Materials and Methods

### Cell culture

Human Cervical Keratinocytes (HCKs) were received from Dr. Aloysius Klingelhutz (donor culture C415, CX2399, CIN612-9E, CIN612-6E). Human Foreskin Keratinocytes (HFKs) were isolated from donor tissue in accordance with University of Arizona IRB. Keratinocytes were passaged in Rheinwald-Green medium (3:1 Ham’s F12 (GIBCO 11765-054) / 4.5 µg/ml glucose Dulbecco’s modified Eagle’s medium (GIBCO 11960-044), 5% fetal bovine serum (Millipore Sigma F8067-500ml lot #16A328), 0.4 µg/ml hydrocortisone (Millipore Sigma H-4001), 8.4 ng/ml cholera toxin (Millipore Sigma 227036), 24 µg/ml adenine (Millipore Sigma A-2786), 6 µg/ml insulin (Millipore Sigma I1882), 10ng/ml epidermal growth factor (Thermo PHG0311), 4 mM L-glutamine (Thermo 25030-149), 50 µg/ml Penicillin/Streptomycin (Thermo 15140-148)). Keratinocytes were incubated with 10% CO_2_. All other cell lines were incubated with 5% CO_2_. For routine passaging F media was changed every 48hrs. For PRMT inhibitor and doxycycline shRNA knockdown experiments F media was changed every 24hrs. Rheinwald-Green media was supplemented with Y-27632 (Chemdea CD0141) at a concentration of 10 µM for passaging. Y-27632 was removed from the media before experiments. Keratinocytes were grown on a layer of lethally irradiated J2-3T3 murine fibroblasts. J2-3T3s were passaged in DMEM supplemented with 10% Neonatal Calf Serum (Thermo 26010-066), 2 mM L-glutamine, and 50 µg/ml Penicillin/Streptomycin. One day before keratinocyte splitting 6X10^5 J2-3T3s were plated and allowed to grow overnight. The J2-3T3s were irradiated with 6000 Gray the day of keratinocyte splitting.

HEK293TT cells were a kind gift of J.T. Schiller (NIH, Bethesda, USA). The HEK293T cells were a kind gift of Samuel Campos (University of Arizona, Tucson, USA). hTERT-immortalized neonatal human foreskin fibroblasts were a kind gift of Felicia Goodrum (University of Arizona, Tucson, USA). 293T, 293TT, and hTERT fibroblasts cell lines were cultured in DMEM supplemented with 10% Fetal Bovine Serum, 2mM L-glutamine, and 50 µg/ml Penicillin/Streptomycin.

### Generation of PRMT1 shRNA cell lines

Primary HCKs were transduced with a pool of 3 doxycycline inducible shRNA lentiviral vectors (Dharmacon RHS4696-200762374, RHS4696-200762016, RHS4696-200687101). Transduced HCKs were treated for 3 days with 640ng/ml doxycycline and TurboRFP expressing cells were sorted using a BD FACS Aria II.

### Chemicals and small molecule inhibitors

The type 1 PRMT inhibitor GSK3368715 (PRMTi) (Cayman chemical, 34886; 1mM stock in water) was used at a final concentration of 2 µM. During PRMTi treatment media was changed every 24hrs for PRMTi treated cells and vehicle (water) control cells. Doxycycline hyclate (Millipore Sigma, D5207; 50 mg/ml in water) was used at a final concentration of 640ng/ml in F media. During doxycycline treatment media was changed every 24hrs for doxycycline treated cells and vehicle (water) control cells. Y-27632 (Chemdea, CD141; 10mM stock in water). Y-27632 was used at a final concentration of 10µM in F media. Media containing Y-27632 was changed every 48hrs. Puromycin (Thermo, J61278.MB; 10 mg/ml stock in water) was used at a final concentration of 300 µg/ml in F media.

### HPV quasivirus and HPV pseudovirus preparation

HPV18 genomes were previously described^43^. 293TT cells were transfected with 15µg of recircularized HPV18 genome, 15 µg of HPV16 pXULL L1/L2 packaging plasmids, 5µg pMEP HPV18 E1, 5µg pMEP HPV18 E2. 48hrs later cells were collected and lysed in 9.5 mM MgCl2 Dulbecco’s Phosphate Buffered Saline (DPBS), 0.35% Brij58, 25 mM ammonium sulfate, 2% RNace-IT cocktail and incubated for 20-24hrs at 37^C^. Next cell debris was removed from solution by centrifugation at 5000g, 5min, at 4^C^. To enhance virion recovery the cell pellet was flash frozen in liquid nitrogen and washed with DPBS. 20% 0.8M NaCl DPBS was added, and the supernatant was loaded onto a CsCl gradient (1.25 g/ml CsCl light 1.4g/ml heavy). Gradients were centrifuged for 16-24 hrs at 20,000 RPM at 18^C^. Quasivirions were collected and washed with Virus Storage Buffer (25mM HEPES ph 7.5, 500mM NaCl, 1mM MgCl2) using 100kDa vivaspin concentrators. Quasivirus was stored at −80^C^ for long term storage.

Viral aliquots were thawed and digested using the turboDNAse kit (Thermo AM1907). 5 µl of the DNAse treated virion solution was then digested by incubating the solution with 95 µl 20mM Tris pH 8, 20mM DTT, 20mM EDTA, 0.5% SDS, 0.2% proteinase K for 20 minutes at 50^C^. Viral DNA was recovered using Zymo DNA Clean and Concentrate kit (Zymo, D4003). Viral genomes equivalents (VGE) per µl were quantified by qPCR.

### shRNA Lentivirus preparation

293T were transfected via PEI-glucose with 14.6µg of packaging plasmids psPAX2 (addgene 12260), 7.9µg of pMD2.G (addgene 12259), and 5ug each of the following PRMT1 targeting shRNA vectors RHS4696-200762374, RHS4696-200762016, or RHS4696-200687101 (Dharmacon). 24hrs post transfection media was changed to F media. Viral supernatant was harvested at 48hrs post transfection, filtered through a 45µM filter, supplemented with 10 µM Y-27632 and 10 µg/ml polybrene. J2s were removed using 0.48 mM EDTA in DPBS 2ml of viral supernatant with 10 µM Y27632 10 µg/ml polybrene was added to each well of keratinocytes. The cells were incubated with viral supernatant for 16hrs. After 16hrs viral supernatant was removed and replaced with F media + Y27632.

### Primary keratinocyte infection assay

6X10^5 keratinocytes were plated per well of a 12 well plate in F media containing 10 µM Y-27632. 24hrs later the media was changed to F media without Y-27632. The next day (Day of infection) J2s were removed. HCKs were allowed to recover in F media for 6 hrs at 37^C^. Following this recovery phase, HPV18 quasivirus was added at a 100 VGE/cell. For mock infections an equivalent volume of F media mixed with Virus Storage Buffer was added. 16 hrs post infection irradiated J2s in F media were added back to the plate.

### qPCR reactions

DNA was then extracted using the Qiagen DNeasy Kit (Qiagen 69504). 1 ng of a reference plasmid pGL3 (Promega U47296) was spiked into the DNA samples prior to loading on to the spin columns to normalize DNA elution efficiency.

RNA extraction was run as per the RNeasy Kit protocol. Residual DNA was digested from RNA samples using TurboDNAse (Thermo Fisher Scientific AM1907). 1 µg of RNA was reverse transcribed using Superscript IV First-Strand Synthesis System and its Oligo dT primer (Thermo Fisher Scientific 18091200).

qPCR was run using PowerUP SYBR Green Master Mix (Thermo Fisher Scientific A25742) and 12.5ng of DNA or cDNA template per 10ul total reaction.

### Western blot

Prior to protein harvest J2-3T3 fibroblasts (J2s) were removed. 5X10^5 keratinocytes were lysed in 200µl of RIPA lysis buffer (50 mM Tris-HCl pH 8.0, 150 mM NaCl, 1% NP40, 0.5% sodium deoxycholate, 1% SDS) with 1x protease inhibitor cocktail (Sigma P1860) and 1µM PMSF (Thermo Fisher Scientific 36978). Total protein concentration was measured with the Pierce BCA Protein Assay Kit (Thermo Fisher Scientific 23227). Rabbit monoclonal anti-PRMT1 (1:2000, Millipore 07-404), rabbit monoclonal anti-GAPDH (1:5000, Cell Signaling 14C10), Rabbit monoclonal anti-HPV18 E7 (1:100, Santa Cruz Bio sc-365035), Rabbit monoclonal anti-ADMA (1:1000, Cell Signaling Technology 13522S) were prepared in 50% Intercept PBS Blocking Buffer 50% Tris-buffered saline containing 0.1% Tween (TBST). Primary antibodies were incubated overnight at 4^C^. Blots were imaged with the LICOR Odyssey Infrared Imaging System and band intensities were measured using the LICOR Odyssey software.

### HPV18 URR promoter assay

Cells were treated with 640 ng/ml doxycycline for six days or 2uM PRMT1 inhibitor for 2 days prior to transfection. Keratinocytes were transfected with 825 ng of HPV18 URR pGL4 (Addgene, 22859) and 125 ng of a renilla luciferase control plasmid pcDNA3.1-b2AR-hRLucIII (A kind gift from Nikolaus Heveker, Canadian Institutes of Health Research) per well using 4µl lipofectamine 2000 per 1 µg of DNA. Lipofectamine complexes were added directly to the F media. 24hrs post transfection luciferase signal was quantified using the Dual-Glo Luciferase Assay System with on plate cell lysis (Promega, E2920).

### Organotypic raft cultures

Organotypic raft cultures were generated as described. Briefly, hTERT-immortalized neonatal human foreskin fibroblasts were embedded in rat tail type I collagen (C3867) to create dermal equivalents in 6 well plates and incubated overnight in 10% FBS DMEM. 24hrs later primary keratinocytes were seeded at 2.5X10^5 cells per dermal equivalent in F. 48hrs after keratinocytes were plated dermal equivalents were lifted onto a hydrophilic 0.4 µm pore size polytetrafluoroethylene membrane to generate an air liquid interface. From the day of lifting to the day of harvest (15 days post lift) media was switched to differentiation media (F media without EGF and supplemented with 1.88 mM CaCl2). Differentiation media was changed every 2 days.

### *In situ* mRNA detection and quantification

PRMT1 mRNAs were detected in formalin-fixed paraffin-embedded 3D organotypic rafts using the RNAscope® Multiplex Fluorescent Detection Kit v2 (ACD, Cat No. 323110) as per manufacturer instructions. A custom probe was designed to target human PRMT1 (ACD, Probe ID. Hs-PRMT1-O1-C2) and labelled using 570 TSA Vivid Dye. DAPI was used as a nuclear stain. Five non-overlapping fields of view were captured for each sample using a Nikon AX R Laser-Scanning confocal microscope (UACC Shared Microscopy Resource). The number of PRMT1 foci per cell were quantified using a custom pipeline including ImageJ/Fiji (PMID: 22743772) and CellProfiler (PMID: 17076895). Briefly, nuclear (DAPI) and PRMT1 channels were split to individual images using ImageJ/Fiji. Nuclei were identified using CellProfiler and expanded to approximate cell segmentation. PRMT1 foci were identified and quantified for each cell.

### Colony forming assay

The colony forming assays were performed as described. 640 ng/ml doxycycline was added to cells 3 days before infection. Two days before infection, 4X10^5 keratinocytes were seeded onto a layer of 1X10^6 irradiated J2-3T3 fibroblasts (J2s) in a 10 cm dish. Y-27632 was removed from the media 24hrs prior to infection. Cells were infected using 100vge. G418 selection was started 2 days post infection at 100 µg/ml (Thermo Fisher Scientific J63871.AB). At 6 days post infection, G418 concentration was lowered to 50 µg/ml. G418 was maintained at 50 µg/ml for the rest of the experiment. Doxycycline was removed at 7 days post infection. HCK colonies were allowed to grow to macroscopic size at which point they were stained with 2.5 mg/ml MTT in F media for 4 hours. Cells were washed with DPBS and imaged under white light on a BioRad Gel Doc system. After imaging, cells were incubated at 37^C^ in DMSO to solubilize the intracellular formazan. Solubilized formazan was quantified using 590 nm absorbance using a DTX-800 multimode plate reader (Beckman Coulter).

### Keratinocyte growth curves

Cells were treated with doxycycline as described for the colony forming assay. However, cells were not infected nor treated with G418. Keratinocytes were serially passaged and counted using a Countess automated cell counter (Thermo) every 4 to 5 days, for a total of 27 to 31 days. Population doubling was calculated using the equation:

population doublings = log 10 (N/No) * 3.32; where N = total number of cells on the 10cm dish on day of serial passage and No = number of cells at time of seeding the 10cm dish.

### Nonsense mediated decay inhibition assay

6X10^5 keratinocytes were plated onto a 12 well plate with 1X10^6 J2 fibroblasts. Cells were treated with PRMTi (2uM) or water vehicle control for 72 hrs before puromycin addition. Cells were treated for an additional 8 hours with water only vehicle control, 2uM PRMTi in F media, 300 µg/ml puromycin in F media, or 2uM PRMTi + 300 µg/ml puromycin in F media for 8hrs. Total cellular RNA was harvested, reverse transcribed, and measured by qPCR as described above.

## Acknowledgements

None.

## Funding

Koenraad Van Doorslaer: HHS NIAID R01AI165638 - 01A1

HHS NIAID: R21AI173916

State of Arizona Improving Health TRIF

State of Arizona ABRC funding

Robert Jackson: Gouvernement du Canada | Natural Sciences and Engineering Research Council of Canada (NSERC): PDF-546182-2020

Fellowship support from the BIO5 Institute

